# Mechanistic insights into chromatin modulation by the “orphan” remodeler ATRX

**DOI:** 10.64898/2026.01.04.697590

**Authors:** Suling Duan, Tsz-Fung Wong, Xian Yang, Huimin Xi, Yanxiang Zhao, Zhong-Ping Yao, Yuanliang Zhai, Jianyou Liao, Tao Ni, Keda Zhou, Yang Liu

## Abstract

Alpha-thalassemia/mental retardation syndrome X-linked protein (ATRX) is an ATP-dependent chromatin remodeler that performs diverse functions, including spacing nucleosomes, facilitating the deposition of histone variant H3.3, and preventing replication stress. The mechanism by which ATRX modulates chromatin is unknown. Here, our cryo-EM structure revealed that ATRX recognizes and modulates the nucleosome through its C-terminal regions by embracing and unwrapping ∼18 base-pair nucleosomal DNA near the entry/exit sites in the presence of ADP•BeF_3_. Strikingly, the DNA minor groove between SHL6 and SHL7 is profoundly deformed by a “pin-like” structure in ATRX, which uniquely intercalates the aromatic side chain of a phenylalanine into the base stackings. Biochemical assays revealed the crucial role of this phenylalanine in sliding the nucleosome without affecting the ATPase activity. Such a “pin-like” structure mimics a “foot-step” on the DNA ladder, which differentiates ATRX from other remodelers. To translocate the DNA on the nucleosome, two more features, including a disordered insertion that contacts the histone surface and a positively charged loop bridging the cross-gyre interactions, serve as essential anchors for the nucleosome modulation by ATRX. Together, we propose a “climbing” model for how ATRX mobilizes the DNA upon ATP hydrolysis.

## Introduction

In all eukaryotes, genomic DNA is packed into the basic form of nucleosomes, which govern access to the genetic information and shape chromatin structure^1^. Chromatin remodelers are ATP-dependent DNA translocases that mobilize the DNA on nucleosomes through their conserved ATPase motor domains, with other auxiliary domains/subunits regulating their engagement with the chromatin substrates for diverse functions^2^. Alpha-thalassemia/mental retardation syndrome X-linked protein (ATRX) is a well-known but little-characterized remodeler^2^. Mutations in ATRX were reported to cause mental disorders^3^ and glioma^4,5^, highlighting its critical role in neurodevelopment.

ATRX functions at heterochromatin regions such as the telomere and peri-centromere^6,7^. A recent study also highlighted its role in regulatory elements such as rDNA and GC-rich euchromatin regions^8^. The N-terminal regions of ATRX (ATRX-N: ADD+RBR), including the H3K9me3 reader domain ATRX–DMNT3L–DNMT3A (ADD)^9,10^ and RNA binding region (RBR)^11^, determines its genomic localization and increases its local concentration through condensate^12^. The C-terminal regions of ATRX (ATRX-C: DBD+ATPase+CTD), comprising the histone chaperone ‘Death Domain Associated Protein’ DAXX binding domain (DBD)^13–15^, the ATPase motor domain (ATPase) and the C-terminal domain (CTD), carry out remodeling activities to maintain heterochromatin structures (Figure 1A). By recruiting DAXX through the DBD, ATRX facilitates the DAXX-mediated H3.3 deposition^15,16^. The ATPase motor plays a central role in spacing the nucleosomes and eliminating repressive DNA structures^15,17,18^, while the underlying mechanism for ATRX remodeling functions remains elusive. In this study, we performed both biochemical assays and cryo-electron microscopy (cryo-EM) to gain insights into how ATRX recognizes and slides nucleosome substrate.

**Figure 1:**
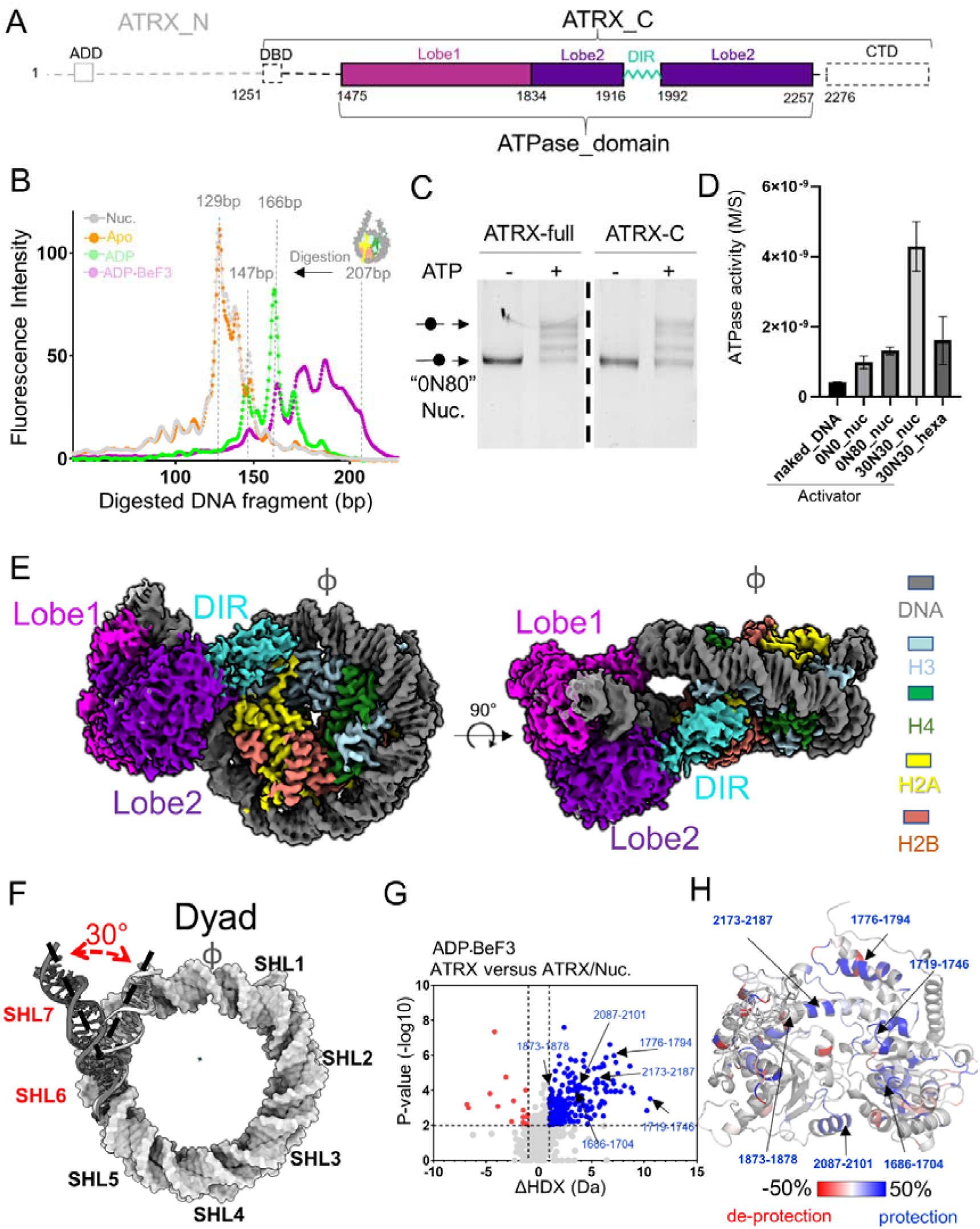
ATRX recognizes and modulates the nucleosome through C-terminal regions. **A.** Domain structure of ATRX. ATRX-N: N-terminal regions; ADD: ATRX–DMNT3L–DNMT3A; ATRX-C: C-terminal regions; DBD: DAXX binding domain; DIR: disordered insertion region; CTD: C-terminal domain; **B.** Micrococcal nuclease (MNase) digestion analysis on the engagement between the nucleosome and ATRX-C in Apo, ADP-bound, and ATP-bound states. The “Nuc.” represented the digested pattern of nucleosome reconstituted on the 207 bp DNA with “601 widom” sequence at the center, and digestion was by 10U MNase for 10 minutes at room temperature. “Apo”, “ADP”, and “ATP” represented the digestion pattern of the nucleosome in complex with ATRX-C in each state under the same digestion conditions. **C.** ATRX-C is sufficient to slide the nucleosome. In the presence of ATP, both full-length ATRX and ATRX-C remodeled nucleosomes with an 80 bp linker on one end (0N80) at a similar level. **D.** The nucleosome with linker DNA on both ends is the best activator for the ATRX ATPase function. The naked DNA (147 bp), nucleosomes (without linker DNA “0N0”, with 80bp linker on one end “0N80”, with 30bp linker on both ends “30N30”) and hexasome (“30N30” DNA template) were tested in the ATPase activity assay. **E.** The overall structure of ATRX-C bound to a nucleosome. The components of a nucleosome were colored as indicated, and the ATRX was colored according to the colors in A; **F.** The DNA between SHL6 and SHL7 is unwrapped by about 30°. **G.** The HDX difference between ATRX-C and ATRX-C with nucleosome substrates in the presence of ADP•BeF_3_. Volcano plot compares the average HDX of ATRX-C and ATRX-C with nucleosome substrates, including all exchange differences at 3600 s. The blue and red dots show significant cut-offs of Δ average |HDX|>0.994 Da and p-value<0.01 from Welch’s t-test. Peptides of ATRX-C with decreased deuterium uptake in complex form are labeled in blue, while the increased peptides are labeled in red. **H.** The ATRX^ATPase^ is colored based on significant changes in deuterium uptake, as shown in G. Colors were generated by the level of HDX changes (normalized in the scale from -0.5 to 0.5 with color key red-white-blue, done in PyMOL). The peptides in the ATRX^ATPase^ that contact the nucleosome are indicated in the volcano plot as well as in the structure model.

## Results

### ATRX remodels the chromatin through its C-terminal regions containing the ATPase domain

Earlier DNase I digestion assay revealed that ATRX stably engages with the nucleosome at the entry/exit site, mainly in the presence of ATP^14^. We therefore hypothesized that the C-terminal regions surrounding the ATPase domain (ATRX-C: DBD+ATPase+CTD) play the central role in nucleosome recognition and modulation. To characterize how ATRX-C engages with the nucleosome, we applied the Micrococcal nuclease (MNase) digestion assay. Consistent with the DNase I digestion results of full-length ATRX, our results demonstrate that the ATRX-C vigorously protected the DNA at the entry/exit sites of the nucleosome only in the presence of ATP analogue ADP•BeF_3_ (Figure 1B and Figure S1). Using the *in vitro* remodeling/sliding assay, we validated that ATRX-C is sufficient to slide the nucleosome on its own to a similar extent as full-length ATRX (Figure 1C). Compared to naked DNA, the nucleosome substrate was a better activator of ATRX-C ATPase activity (Figure 1D). Intriguingly, we revealed that the nucleosome with linker DNA on both sides shows much higher efficiency in stimulating ATPase activity (Figure 1D), suggesting the vital role of extra-nucleosomal DNA for the remodelling activities, as was also observed for other remodelers^19–24^.

Together, we demonstrated that the ATRX-C (DBD+ATPase+CTD) is the main driving force that recognizes and modulates the nucleosome, while ATRX-N (ADD+RBR) may play an auxiliary role in facilitating remodeling. Therefore, our following study focused on ATRX-C to unravel its detailed remodeling mechanism.

### Structural determination of ATRX-nucleosome engagement

To reveal how ATRX recognizes the nucleosome and modulates its structure, we applied cryo-EM to determine the structure of the complex containing ATRX-C in Apo, ADP-bound, and ATP-bound (ADP•BeF_3_) states with mono-nucleosome (30N30, 30 bp DNA on each end). The complexes were reconstituted and stabilized by GraFix (Gradient Fixation), similar to other remodeler-chromatin complexes^24,25^ (Figure S2). The ATRX in the Apo or ADP-bound state was so unstable on the nucleosome that ATRX density was barely visible or recognisable after 3D reconstruction. The ADP•BeF_3_-bound ATRX-C was much more stably associated with the nucleosome core, allowing us to reconstruct its density map (Figure S3 and S4). This aligns with our MNase digestion results (Figure 1B and Figure S1). The association of ADP•BeF_3_-bound ATRX on mono-nucleosome in our structure is highly flexible (Figure S3). Although the DAXX binding domain (DBD) and C-terminal domain (CTD) were included in the construct, only the ATPase domain was resolved at high resolution. This suggests that the ATPase domain is the only stably bound part of ATRX-C on the nucleosome substrate. Through focus refinement, we determined the structure of nucleosome-bound ATRX^ATPase^ at a resolution of 3.4 Å (Figure S4).

### ATRX opens the nucleosome through its ATPase domain

Overall, the ATRX^ATPase^ resembles a “bottle opener” that unwraps the DNA at the location between Super Helical Locations (SHL) 6 and 7, which is at the entry or exit of the nucleosome (Figure 1E). Utilizing the histone surface and the second DNA gyre as fulcrums, ATRX embraces and unwraps >18 base-pairs (bp) of DNA from the histone core, thereby disrupting key histone-DNA interactions made by H3 □N and the H2A loop 2 (L2) (Figure 1E and 1F). This specific engagement is highly dynamic, with ATRX rotating and rocking the DNA to different positions (Figure S3C), providing a unique modulation mode compared to other known remodelers.

To validate our structure in solution, we employed hydrogen/deuterium exchange-mass spectrometry (HDX-MS), by comparing deuterium uptake in ATRX-C and ATRX-C bound to its nucleosome substrate (Figure 1G). Within ATRX-C, a large number of peptides in the ATPase domain showed significantly decreased deuterium uptake upon nucleosome interaction (ΔHDX > 0.994 Da, p-value < 0.01; see methods). The interaction interfaces identified by HDX-MS are consistent with our cryo-EM structure, and peptides facing the nucleosomal DNA were all highly protected from exchange (Figure 1H). To explore the impact of ATRX-C on the nucleosome structure in the ADP•BeF_3_-bound state, we compared the protection of histones in the nucleosome to the histones in the ATRX-nucleosome complex. Most histone residues showed no significant changes in deuterium uptakes, while some histone peptides buried in the nucleosome core (H3: 84-99, H3: 104-110, and H2B: 2-41) showed significantly increased deuterium in the presence of ATRX-C (Figure S5), indicating a destabilizing effect of nucleosome by the engagement of ATRX in the ADP•BeF_3_-bound state.

The high resolution of the ATRX density map allowed us to build an unambiguous atomic model for the majority of the ATPase domain (Figure 2A and Figure S4). Two RecA-like lobes are located on opposite sides of the DNA, with a catalytic pocket hosting a nucleotide close to the junction (Figure 2A). Consistent with other studies on ADP•BeF3-bound remodelers, the ADP density was apparent, while the BeF3 was not visible, which mimics ATP in an activated reaction intermediate state. The ADP•BeF3-bound ATRX ATPase adopts a closed conformation to embrace the DNA. At first glance, the catalytic core of the ATPase around the nucleotide-binding pocket is highly conserved between ATRX and other conventional remodelers, such as Snf2h, with an RMSD (root-mean-square deviation) of the amino acid backbone of less than 1 Å (Figure S6). Key residues that interact with the nucleotide are identical to those of other remodelers (Figure 2A)^19,20,23,24,26^. The brace helix, which was revealed to be essential for coupling ATP hydrolysis to DNA translocation^25^, is located on top of the guide strand in the same manner as ISWI remodelers (Figure 2A). While the ATPase core of ATRX shares high similarity with other remodelers, additional elements surrounding the ATPase core define its unique features in its engagement with the nucleosome.

**Figure 2:**
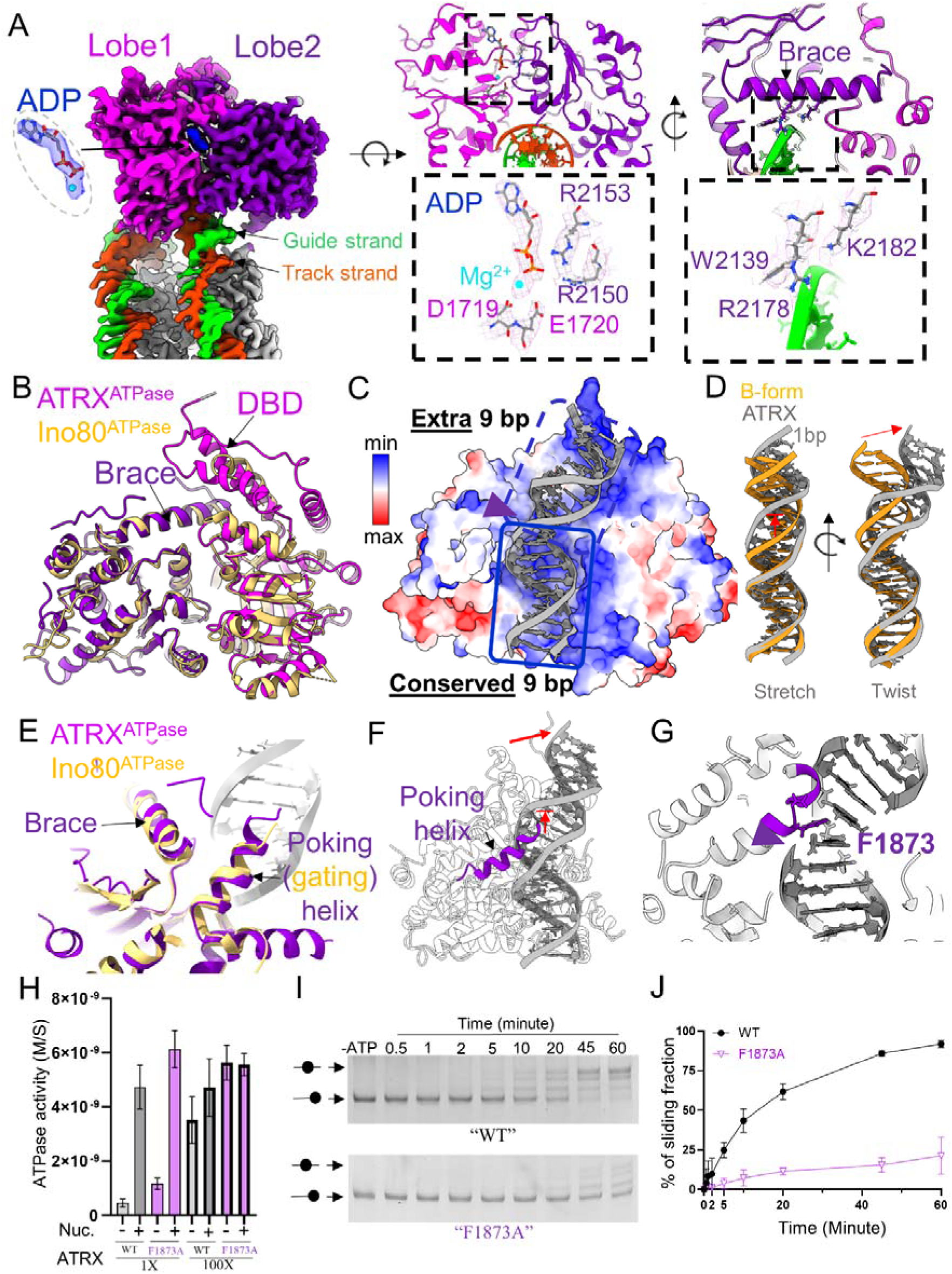
ATRX^ATPase^ deforms the DNA through a “pin-like” structure. **A.** The ADP•BeF_3_-bound ATRX^ATPase^ embraces DNA between SHL6 and SHL7. The individual lobes are colored as indicated. The density of “ADP” with “Mg2+” was clearly visible in the ATPase pocket. The key amino acids around the nucleotide were conserved compared to other remodelers. The “Brace” helix interacts with the guide strand through conserved amino acids as well. **B.** ATRX^ATPase^ contains an additional DNA-binding domain (DBD) compared to Ino80^ATPase^ (pdb:6hts). The ATPase domains were superimposed (yellow represents Ino80, and pink and purple represent the two lobes of the ATRX ATPase). **C.** The electrostatic potential of the surface on the ATRX^ATPase^ directly contacts the DNA. “Conserved 9bp” and “extra 9bp” were based on a comparison with the Snf2h/DNA contacts (pdb 8v4y); **D.** ATRX^ATPase^ deforms the contacting DNA so that the minor groove is significantly widened, causing a 1-bp stretch and twist of the DNA. **E.** The □15 helix (often called the gating helix in other remodelers) of ATRX^ATPase^ pokes into the minor groove of the DNA. **F.** The poking helix pushes the guide strand to widen the DNA groove. **G.** A phenylalanine (F1873) at the tip of the “poking helix” is inserted into the minor groove to form π-π interaction with DNA bases in the guide strand. **H.** “Pin” removal mutation (F1873A) does not affect the nucleosome-dependent ATP hydrolysis. The ATPase activities of wild-type ATRX-C and the mutant were measured in the absence or presence of the nucleosome substrate (30N30) at the indicated ratios. **I.** “Pin” removal mutation (F1873A) dramatically impairs the nucleosome sliding function of ATRX-C. **J.** Quantification of the sliding assay from three biological replicates of F1873A.

### Prolonged DNA interaction through an extra DNA-binding domain in ATRX^ATPase^

The first distinct feature of ATRX is its interaction with nucleosomal DNA. Typically, Snf2 ATPase domains directly contact ∼8-10 bp of double-stranded DNA, mainly along one strand^19,24–28^. In ADP•BeF_3_-bound ATRX^ATPase^, the positively charged surface embraces a two-fold longer DNA segment than Snf2h^ATPase^ (Figure 2B and 2C). The conserved catalytic core of ATRX^ATPase^ contacts ∼9 bp of DNA, and ATRX^ATPase^ forms a second DNA-binding domain that grabs an additional ∼9 bp of DNA. This results in at least 18 bp nucleosomal DNA being bound and unwrapped from the histone core by ATRX (Figure 2C). Since nucleosomal DNA is highly curved every 10 bp periodically, ATRX^ATPase^ could not embrace 18 bp of bent DNA inside the nucleosome core particle. Instead, only the DNA at the entry/exit of the nucleosome (SHL6 and SHL7) would allow such interaction through unwrapping.

### DNA deformation by a unique “pin-like” structure in ATRX^ATPase^

More intriguingly, the DNA minor groove is significantly widened and deformed between SHL6 and SHL7, compared to the standard B-form DNA (Figure 2C and 2D). The ATRX □15 (detailed sequence in Figure S7 and S8), often referred to as the “gating helix” in other remodelers, pokes into the DNA groove (referred here as “poking helix”) and lifts one strand of DNA (non-translocation strand), resulting in twisting and 1-bp stretching (Figure 2D and 2E). The lifted and tilted DNA is perfectly positioned to be bound by the second DNA-binding domain of ATRX ATPase (Figure 2B and 2C). R2178 in brace (□22) helix closely contacts the DNA backbone of the lifted strand above □15, which may stabilize the minor groove widening (Figure 2A). Strikingly, phenylalanine F1873 sits at the tip of the □15 helix, and is intercalated into the DNA to form π-π interactions (Figure 2F-G). This “pin-like” mechanism, in which a “wedge” or “pin” like element with a translocase motor domain separates the DNA^29^, is often observed in DNA helicases but was never found in chromatin remodelers^2^, indicating that ATRX might modulate the DNA structure in a way similar to the true DNA helicase. To test whether such a unique “pin-like” feature in ATRX is required for its functions, we removed the tip of the “pin” by mutating the phenylalanine to alanine (F1873A). In the ATPase activity assay, the F1873A mutant showed almost the same level of ATP hydrolysis efficiency as the wild-type ATRX (Figure 2H). However, nucleosome sliding efficiency was dramatically impaired by the F1873A mutation (Figure 2I and 2J), demonstrating that the “pin-like” structure in ATRX is essential for translocating DNA on the nucleosome, a property no other remodelers exhibit.

### Histone contact through a disordered insertion region

Direct histone contact is a common feature in determining the specificity of nucleosome-remodeler interactions^19,20,24,28,30,31^. In our cryo-EM map of the ATRX-nucleosome complex, a density extruding from lobe 2 of the ATPase contacts the histone surface, extensively interacting with the H2A L2 loop, H3 □-N, and H2A C-terminal domain (Figure 3A). The unique disordered insertion region DIR (1905-2000 a.a.) within ATRX lobe2 could fit into this density. Although a longer insertion has been found in the same location in the INO80^ATPase^ (Figure 3A), it is responsible for recruiting the necessary co-factors such as Rvb1/Rvb2 AAA+ ATPase in INO80 complex^19,24^. Here, the ATRX DIR directly serves as one fulcrum on the nucleosome to promote the proper configuration of ATRX on nucleosomal DNA, with a function distinct from that of other remodelers. To test whether the DIR-histone contact is crucial for the ATRX function, we removed the majority of the residues in this disordered region (deletion of 1916 to 1995, ΔDIR). The nucleosome-stimulated ATPase activity of the ΔDIR mutant was significantly lower than that of the wild-type ATRX (Figure 3B). Consistently, we observed reduced nucleosome sliding efficiency in the ΔDIR mutant (Figure 3C-3D). These results suggested that histone contact via the DIR is essential for maintaining the ATRX configuration on the nucleosome, which is required to stimulate ATP hydrolysis for the nucleosome sliding function.

**Figure 3:**
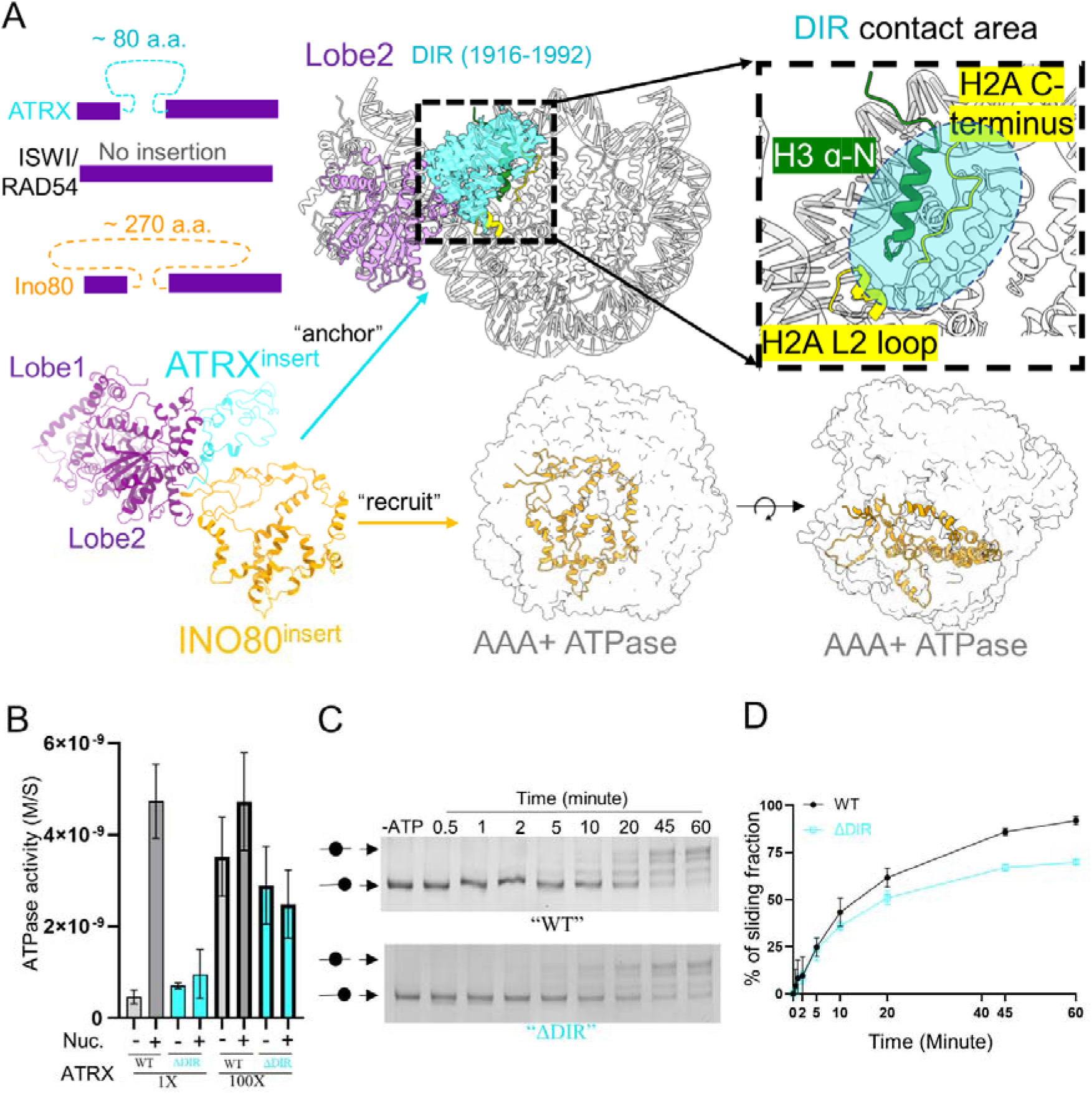
A disordered insertion region supports the nucleosome recognition and modulation of ATRX. **A.** The disordered insertion region (DIR) in ATRX forms extensive contacts with histones. Compared to other remodellers, ATRX and Ino80 ATPase contain long and disordered insertions. ATRX DIR density covers the histone H3 □-N helix, H2A C-terminus, and H2A L2 loop (up-panel). In contrast, Ino80 DIR interacts with other co-factors in the Ino80 complex. **B.** ATRX DIR removal mutation (ΔDIR) significantly reduces the nucleosome-dependent ATP hydrolysis. The ATPase activities of wild-type ATRX-C and the mutant were measured in the absence or presence of the nucleosome substrate (30N30) at the indicated ratios. **C.** DIR removal mutation (ΔDIR) noticeably decreases the nucleosome sliding efficiency of ATRX-C. **D.** Quantification of the sliding assay from three biological replicates of ΔDIR.

### “Cross-gyre” interaction through a positively charged loop

The ATPase of chromatin remodelers often interacts with DNA on the second gyre. Such a “cross-gyre” interaction was also observed in ATRX-nucleosome engagement in the presence of ADP•BeF_3_ (Figure 4A). The 30° unwrapping of the DNA end allows ATRX to approach the nucleosomal DNA at the second gyre, which is stabilized by a positively charged loop (aa 1694-1705, Figure 4A) in ATPase, mimicking a “buckling” action. The close distance positions the side chains of R1698 and K1699 to grab the phosphate backbone of the nearby DNA strand. This interaction could further stabilize the orientation of ATRX on the primary DNA between SHL6 and SHL7. By sequence alignment among the remodelers, this extra loop is unique in ATRX. When the DNA end of the nucleosome is unwrapped, the distance between two gyres increases. Typical Snf2 ATPases could not bridge two gyres in such a configuration on their own (Figure 4A). The ATPase domain in INO80 requires the Les2 helix to interact with the second gyre, while other ATPases only bridge the gyres over a shorter distance (Figure 4A). If the ATRX is superimposed in SHL2 position like Snf2h, the extra loop unique in ATRX will cause a clash with the second gyre (Figure 4B), indicating that SHL6 is the favoured engagement site on the nucleosome for ATRX.

**Figure 4:**
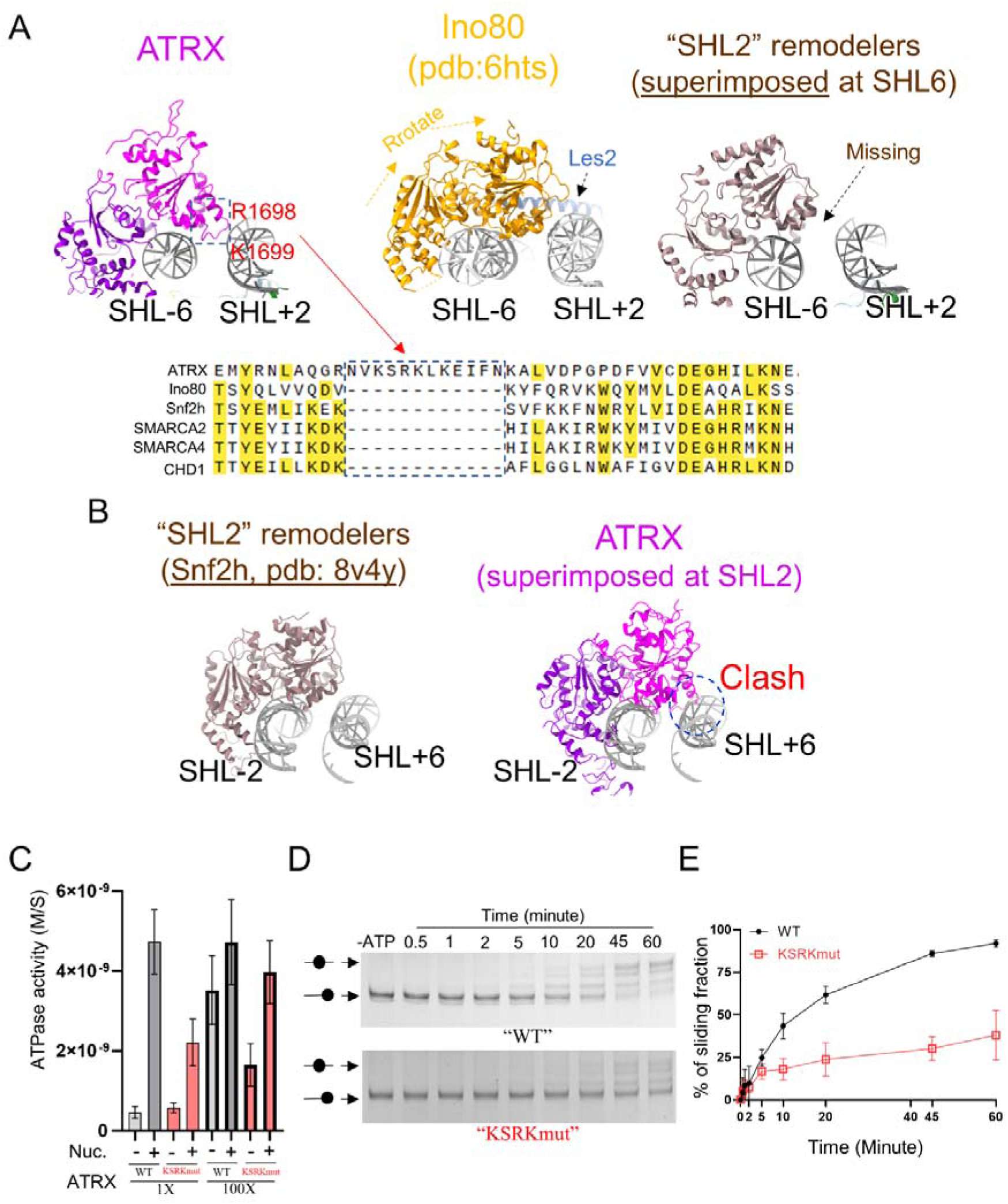
A unique loop of ATRX bridges the “cross-gyre” interaction on the nucleosome. **A**. The positively charged loop for cross-gyre contact is unique to ATRX. Left: ATRX in the complex; Middle: Ino80 ATPase is facilitated by Les2 module to form the “cross-gyre” interaction; Right. Snf2h cannot interact with the DNA on the other gyre at the same position where ATRX engages (superimposed with ATRX). The lower panel is the sequence alignment of ATRX to other well-known remodelers, focusing on the region close to the positively charged loop in ATRX^ATPase^. Only ATRX evolves this extra loop. The red arrows highlight three positively charged amino acids. **B.** The loop in ATRX for “cross-gyre” interaction would cause a major clash with the second gyre if ATRX^ATPase^ is positioned in SHL-2, where most remodelers bind. **C.** Loop mutation (neutralizing the positively charged residues, KSRK mutant) reduces the nucleosome-dependent ATP hydrolysis. The ATPase activities of wild-type ATRX-C and the mutant were measured in the absence or presence of the nucleosome substrate (30N30) at the indicated ratios. **D.** Loop mutation (KSRKmut) decreases the nucleosome sliding efficiency of ATRX-C. **E.** Quantification of the sliding assay from three biological replicates of KSRKmut.

To test whether the “cross-gyre” interaction is crucial for ATRX function, we mutated three positively charged residues in this loop to alanines (KSRKmut, 1696-1699). The nucleosome-stimulated ATPase activity of the KSRKmut dropped less than one fold (Figure 4C). However, we observed a more dramatic inhibition of nucleosome sliding by the “KSRKmut” mutation (Figure 4D and 4E) than by the DIR mutant. This demonstrates that the positively charged loop is essential not only for simulating ATRX ATPase activity but also for promoting DNA translocation along the nucleosome.

## Discussion

### “Orphan” chromatin remodeler ATRX

An early phylogenetic analysis of the helicase region indicated that ATRX is distinct from conventional chromatin remodelers^32^, while the details were missing. Our cryo-EM structure, together with biochemical and HDX-MS data, unravelled how the unique features/elements of ATRX are specifically adapted to recognize and modulate the nucleosome, in a manner different from that of other known remodelers.

Up-to-date, most chromatin remodelers dock their ATPase on the nucleosome at the SHL2 position. However, ATRX ATPase favors SHL6 on the nucleosome, which is determined by three key features: 1) ∼18 bp DNA contact; 2) a uniquely evolved loop for “cross-gyre” interaction; 3) histone contact through DIR. The prolonged DNA contact could not be supported at the SHL2, where DNA is highly curved in the nucleosome. Meanwhile, the positively charged loop responsible for the “cross-gyre” interaction could not engage at the SHL2 without any clash (Figure 4B). Therefore, the DNA ends at nucleosome entry/exit sites are the primary sites where ATRX recognizes and modulates chromatin. This is consistent with earlier DNase I footprint analysis^14^ and our MNase digestion assay. Among other chromatin remodelers, Snf2 was also reported to bind the DNA ends, but without unwrapping the DNA^25^ (Figure S9A). The Ino80 complex is the only one that contacts SHL6 to unwrap the nucleosomal DNA^19,22,24^. However, co-factors are required to assist the Ino80 ATPase in DNA unwrapping (Figure S9A). Therefore, ATRX is by far the only chromatin remodeler that modulates the nucleosome near the entry/exit sites merely on its own (Figure S9).

More strikingly, ATRX evolves a “pin” element on the tip of the gating helix for its DNA translocation along the histone core. The gating helix itself is crucial to couple the ATP hydrolysis to the DNA translocation by regulating the width of the DNA minor groove^33^. In ATRX, the gating helix is further modified to poke into the DNA bases on the guide strand via an aromatic residue (F1873), which induces a 1-bp twist in the DNA. Such DNA intercalation and “pin-like” structure have not been observed in any other chromatin remodelers. Interestingly, this “pin” structure is a classic element of true DNA helicases, which facilitate strand replacement and duplex melting. Since ATRX has been reported to dissolve abnormal DNA structures, such as the G-quadruplex^17,34^, it is reasonable to propose that the “pin-like” structure on the gating helix plays a central role in such functions of ATRX.

### Sliding through “climbing”: ATRX translocates the DNA on the nucleosome

A complete cycle of chromatin remodeling by ISWI during ATP hydrolysis was recently reported^23^. When comparing the ADP•BeF_3_-bound ATRX^ATPase^ to ATP-bound state of ISWI^ATPase^, the catalytic core in ATRX^ATPase^ surrounding the ATP pocket is almost identical to that of ISWI^ATPase^ (Figure 5A). The unique elements we discovered in ATRX are all located at the periphery of the central folding (Figure S6). Therefore, the mechanism of chemo-mechanical coupling upon ATP hydrolysis would be conserved in the ATRX. Based on this, we could model the structures of ADP-bound or apo-state ATRX^ATPase^ to predict ATRX movement along the DNA (Figure 5B): the ATP hydrolysis would trigger a significant conformational rearrangement of ATRX^ATPase^ lobe2, leading to its detachment from the tracking strand (Figure 5C-5D). The gating helix with the “pin” structure in ATRX^ATPase^ will then retract from base stacking. Without the “pin”, the stretched DNA will relax into B-form DNA, causing a 1-bp DNA shift towards the nucleosome dyad (Figure 5E). Such DNA relaxation was also observed in Ino80-nucleosome complexes (from an ATP analogue to ADP^22^). Relatively, with the DNA “stretch to relax”, the “pin” element in ATRX would be positioned 1-bp forward. When ATP binds to ATRX again, the two lobes will adopt the closed conformation, and the gating helix will poke the “pin” element at the new position. In this way, ATRX will move 1-bp in each ATP hydrolysis cycle (Figure 5F). This process resembles “climbing a ladder”: the two lobes of ATRX ATPase are the two hands and the “pin” structure mimics a foot, where the hands and one foot are coordinated to climb the DNA ladder 1-bp per step by consuming ATP. On the nucleosome substrate, the “cross-gyre” interaction and the histone contact serve as two anchors for this climbing, which propels DNA backward. Together, this mode enables ATRX to translocate DNA on chromatin (Figure 5F).

**Figure 5:**
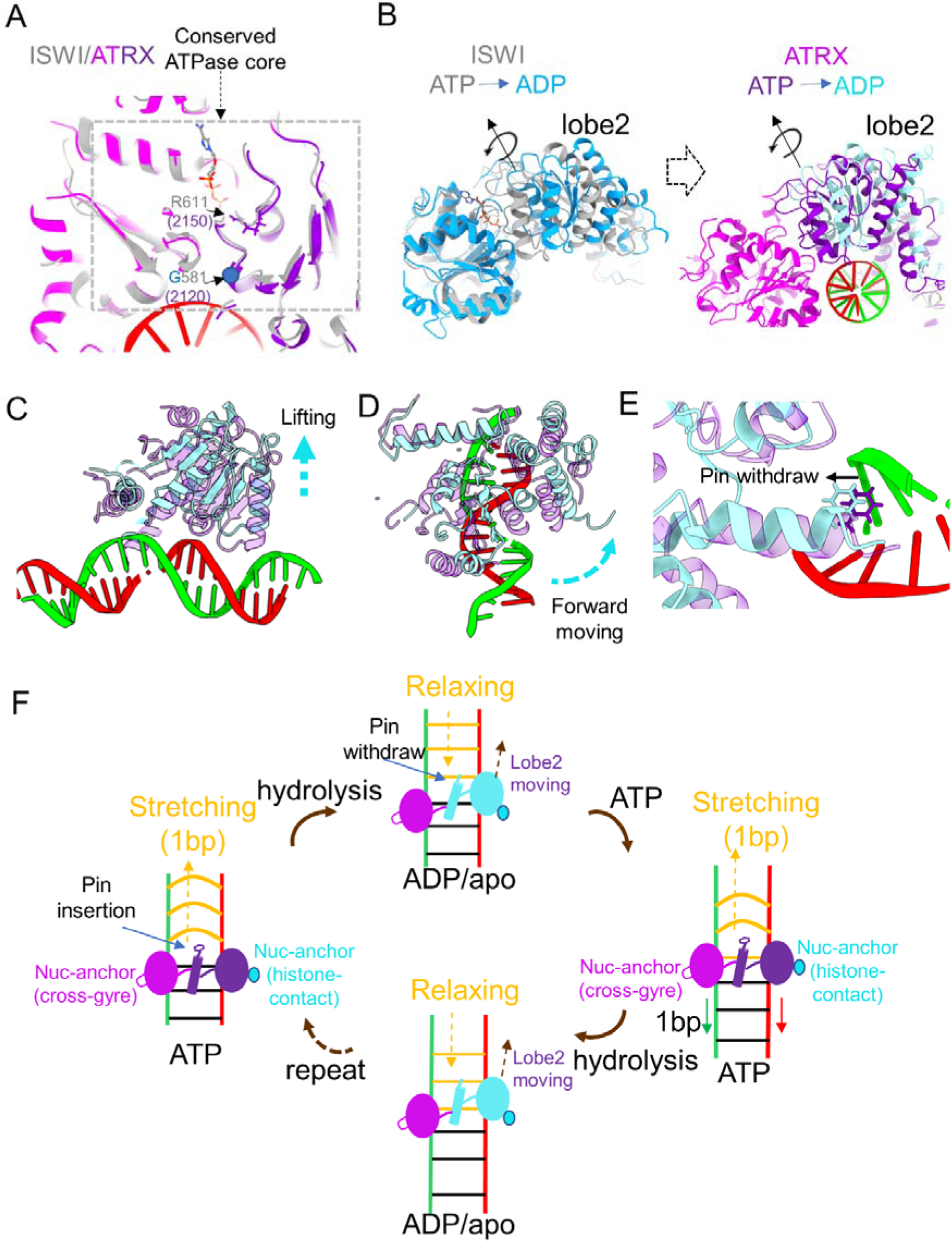
The chromatin remodeling mode for ATRX. **A.** ATRX and ISWI shared an identical ATPase core. ADP•BeF_3_-bound ATRX^ATPase^ was superimposed with ATP-bound ISWI^ATPase^ (pdb: 9jnp). **B.** Simulation of ATRX^ATPase^ conformation change during ATP hydrolysis based on the ISWI data (pdb: 9jnp for ATP-bound, and 9jnu for ADP-bound). **C.** The lobe2 of ATRX^ATPase^ detached from the DNA during the ATP hydrolysis. **D.** The lobe2 of ATRX^ATPase^ moves forward on the tracking strand. **E.** the “pin” on the gating helix of ATRX withdraws from the insertion position at the ADP-bound state. **F.** the cartoon model of “ATRX slides the DNA on nucleosome through climbing”. In the ATP-bound state, ATRX engages the nucleosome at the SHL6-7 to unwrap and dramatically deform the DNA by inserting the pin on the gating helix, mimicking a foot stepping on the DNA ladder. Such an engagement activates ATRX’s ATPase activity, leading to ATP hydrolysis and inducing an open conformation of lobe2. The pin structure is then retracted from the insertion position, causing the DNA relaxation backward. This results in the new position of the gating helix with the pin. When a new ATP binds to the ATPase, the two lobes, mimicking two hands, adopt the closed conformation to grab the DNA strands, and re-insert the pin at the new position, moving the ATPase 1-bp forward. Since the ATPase is anchored by histones and DNA on the second gyre, it would remain in place and propel the DNA backward, which enables the nucleosome sliding along the DNA.

### Disease-related mutations in the ATPase domain

ATRX was well-known for its mutations that cause the Alpha-thalassemia/mental retardation syndrome^3,4,35^. We mapped the reported mutations onto our cryo-EM structure (Figure S10A)^35–37^, and most are buried within the folding regions of the two lobes, suggesting an impact on protein conformation or stability. The L1746S mutation was shown to decouple ATPase activity from DNA activation^36^, thereby disrupting the hydrophobic interaction with V1604 (Figure S10B), leading to a more flexible motif I (P-loop) and potentially initiating ATP hydrolysis prematurely, independent of DNA or nucleosome binding. As suggested earlier, disruption of hydrophobic interactions appears to be a common feature of mutations in the ATPase domain in ATR syndrome^36^. A recently reported mutation Y1758C, which causes neuroinflammation^37^, could disrupt the hydrophobic core to destabilize folding at the end of the “brace” helix (Figure S10C). Similar mutations are found on V1538 and V1552 as well (Figure S10D). Interestingly, multiple aromatic residues were also mutated to cause the ATR syndrome^36^ (Figure S10A). Y1847, which forms a base-stacking interaction with F2210, is crucial for positioning the gating (poking) helix to intercalate the “pin element” (Figure S10E). In addition, charge-charge interactions are essential for folding, and K1650N could reduce its interactions with E1639 and E1643, thereby weakening loop formation (Figure S10F). The side chains of Y2084 and R2085 bridge the DNA-binding loop to the rest of lobe2 via a hydrophobic and a charge-charge interaction, respectively. Mutations in these two amino acids (Y2084H and R2085C) could therefore affect the mechano-compactness (Figure S10G).

The chemo-mechanical coupling for ATPase activation and DNA translocation requires proper folding of the ATPase domain in chromatin remodelers. By far, most mutations found in the ATRX ATPase domain seem to cause local destabilization. Hence, our cryo-EM structure provides a solid foundation for interpreting existing mutations and for evaluating newly emerging mutations for prognostic purposes.

**Table.** Cryo-EM data collection, refinement and validation statistics.

|  | #1 ATRX<br>(Focused map,<br>EMDB-62684) | #2 Nucleosome<br>(consensus map,<br>EMDB-62686) |
| --- | --- | --- |
| (pdb: 9L06, composite model) |  |  |
| **Data collection and processing** |  |  |
| Magnification | 135000 |  |
| Voltage (kV) | 300 |  |
| Electron exposure (e-/Å<sup>2</sup>) | 40 |  |
| Defocus range (μm) | -1.0 to -2.5 |  |
| Pixel size (Å) | 0.9557 |  |
| Initial particle images (no.) | 2,364,881 |  |
| Final particle images (no.) | 147,422 |  |
| Map resolution (Å) | 3.4 | 3.0 |
| FSC threshold | 0.143 | 0.143 |
| Map resolution range (Å) | 3.3-6.0 | 2.9-6.0 |
| **Refinement** |  |  |
| Initial model used (PDB code) | alphafold | 8v4y |
| Model resolution (Å) | 3.6 | 3.1 |
| FSC threshold | 0.5 | 0.5 |
| Model resolution range (Å) | 3.6 | 3.1 |
| Map sharpening *B* factor (Å<sup>2</sup>) | -70.4 | -79.1 |
| Model composition |  |  |
| Non-hydrogen atoms | 6681 | 11362 |
| Protein residues | 753 | 766 |
| Nucleotide | 37 | 260 |
| Ligands | 0 | 0 |
| *B* factors (Å<sup>2</sup>) |  |  |
| Protein | 38.45 | 49.89 |
| Nucleotide | 80.12 | 128.58 |
| R.m.s. deviations |  |  |
| Bond lengths (Å) | 0.003 | 0.013 |
| Bond angles (°) | 0.587 | 1.9 |
| Validation |  |  |
| MolProbity score | 1.75 | 0.5 |
| Clashscore | 3.80 | 0 |
| Poor rotamers (%) | 1.45 | 0.00 |
| Ramachandran plot |  |  |
| Favored (%) | 92.74 | 98.67 |
| Allowed (%) | 7.26 | 1.33 |
| Disallowed (%) | 0 | 0 |

## Acknowledgments

We thank the University Research Facility in Chemical and Environmental Analysis, and the University Research Facility in Life Sciences at The Hong Kong Polytechnic University for their instrumental and technical support. We also thank L.□T.□Fu for assistance with cryo-EM data collection. The cryo-EM grid screening and data collection were performed at the Li Ka Shing cryo-EM laboratory of the Centre for PanorOmic Sciences at The University of Hong Kong. Funding is supported by the Research Grant Council-Early Career Scheme of YL and by the start-up funds of YL and KZ.

## Author contributions

KZ and YL conceived and supervised the project; SD cloned and expressed all the ATRX constructs with the help of YL and HX; SD optimized sample and grid preparation and performed extensive screening with the help of KZ; SD, KZ and YL collected the cryo-EM data. SD, YL, and KZ analyzed the EM data and obtained the electron density maps with the help of TN. KZ modeled and refined the structures. KZ and YL interpreted EM structures and drafted the manuscript. SD and YL designed and performed all biochemical assays, with input from KZ. SD and YL designed and performed the HDX experiments with the help of XY. SD and TW analyzed the HDX data. SD, YL, and KZ wrote the manuscript and prepared the figures. YL, KZ, YXZ, ZPY, YLZ, TN, and JYL edited the manuscript.

## Supplemental Figures

**Figure S1:**
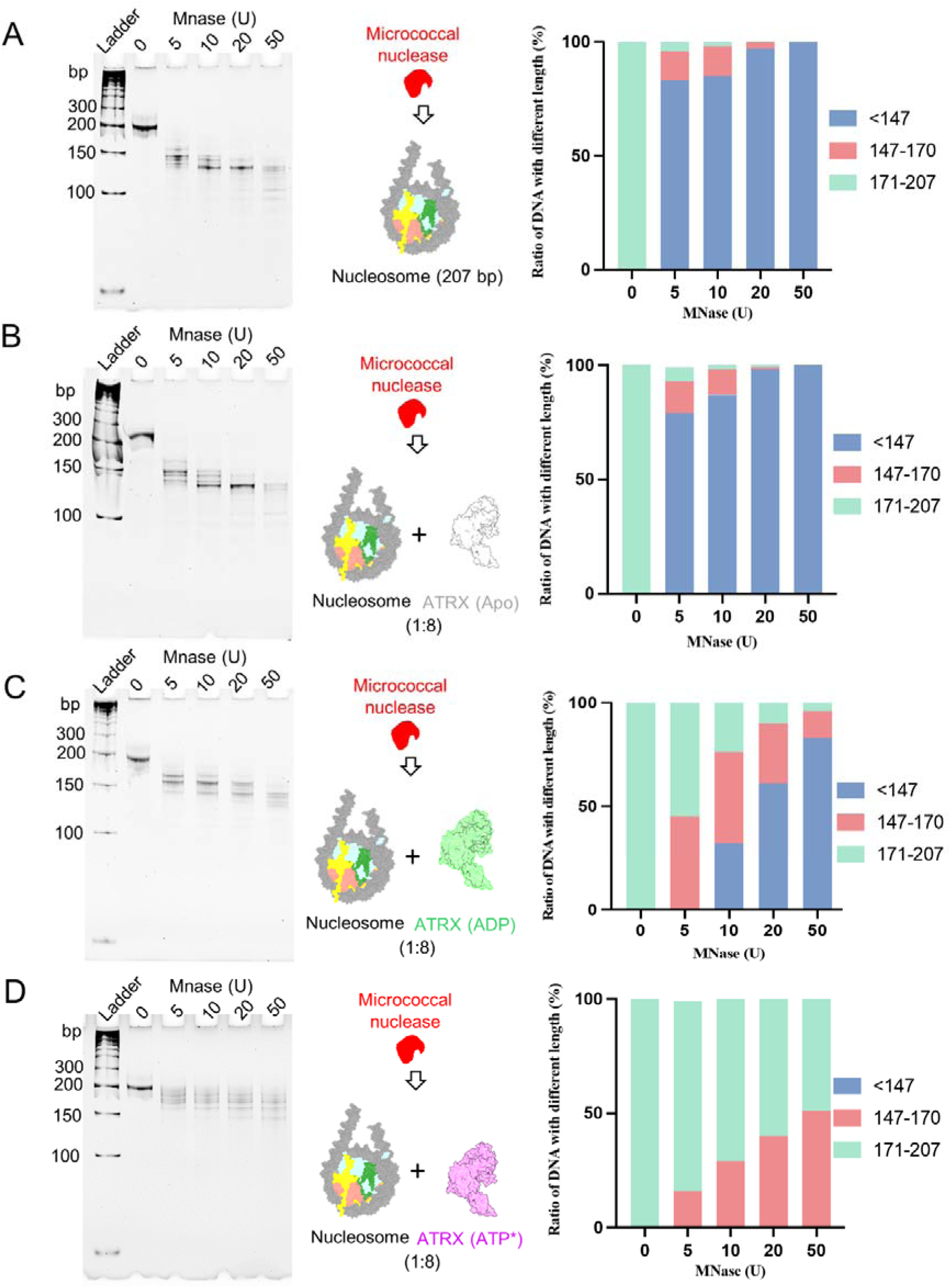
Evaluation of the nucleosomal DNA end accessibility by micrococcal nuclease (MNase) digestion assay. A. MNase digestion results of nucleosome with linker DNA (30 bp at each end). Left, gel-shift assay on native PAGE for resolving the digested DNA fragments from different amounts of MNase. Right, statistics on the length distribution of the digested DNA fragments by bioanalyzer. B. MNase digestion results of Apo ATRX-C in complex with the nucleosome (8:1 ratio in solution). Results are displayed as in a. C. MNase digestion results of ADP-bound ATRX-C in complex with the nucleosome (8:1 ratio in solution). Results are displayed as in a. D. MNase digestion results of ADP•BeF_3_-bound ATRX-C in complex with the nucleosome (8:1 ratio in solution). Results are displayed as in a. ATP* stands for ATP analogue ADP•BeF_3_.

**Figure S2:**
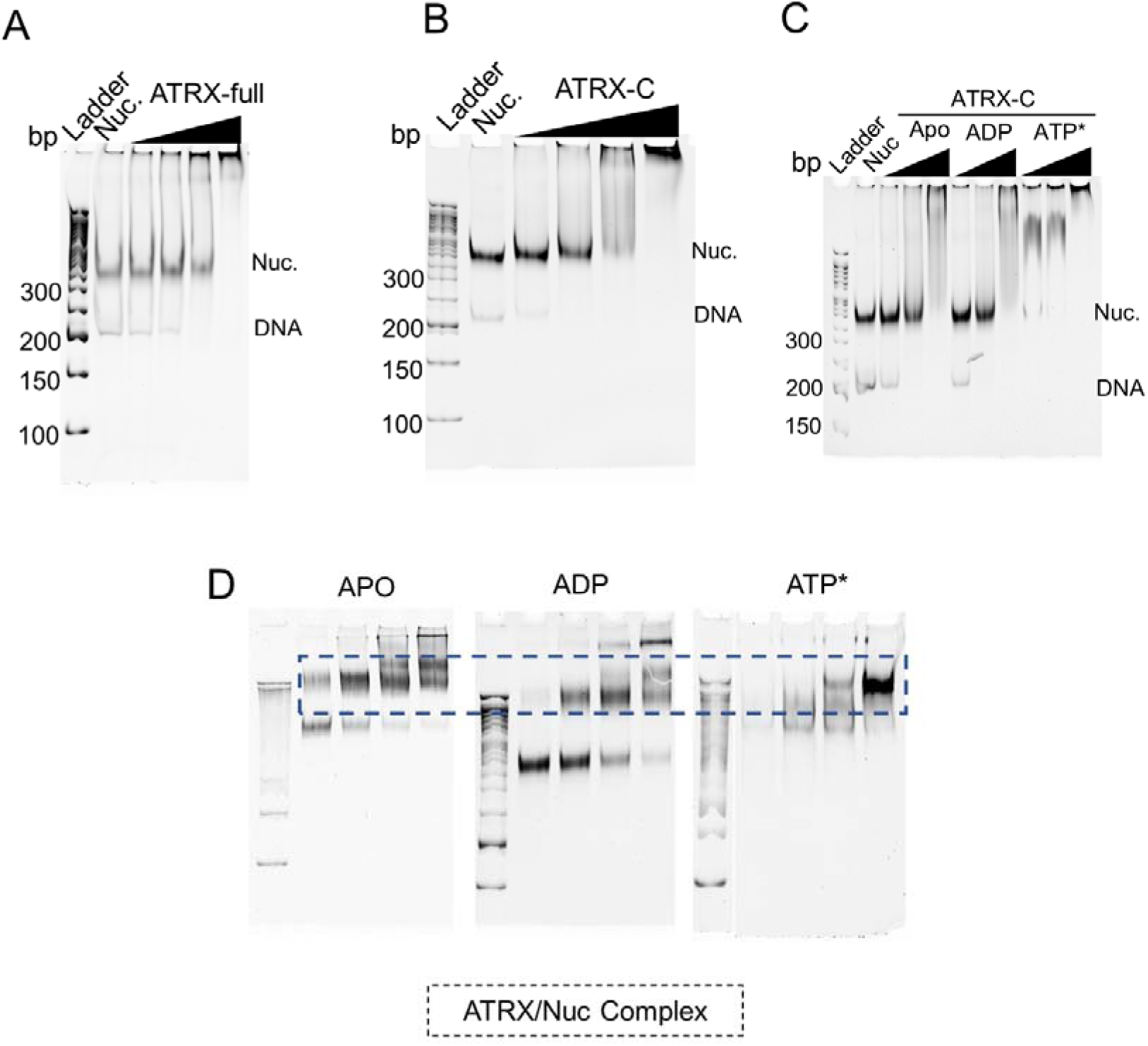
*In vitro* gel-shift assay for ATRX and nucleosome interaction. A. Full-length ATRX was mixed with nucleosome at ratios of: 1:1, 2:1, 4:1, and 8:1, respectively. “Nuc” represents nucleosome reconstituted on the 207 bp “widom 601” DNA sequence. The nucleosome concentration is 250 nM. The native gel-shift was performed on 5% PAGE, and stained by SYBR Gold to indicate nucleosome and complex formation. B. ATRX-C (DBD+ATPase+CTD) was mixed with nucleosome at ratios of 1:1, 2:1, 4:1, and 8:1, respectively. C. The ATRX-C in Apo, ADP-bound, and ATP-bound states interacts with the nucleosome. ATP* represents ATP analogue ADP•BeF_3_. D. The input sample from GraFix for cryo-EM grid preparation contained mainly the ATRX-C/nucleosome complexes.

**Figure S3:**
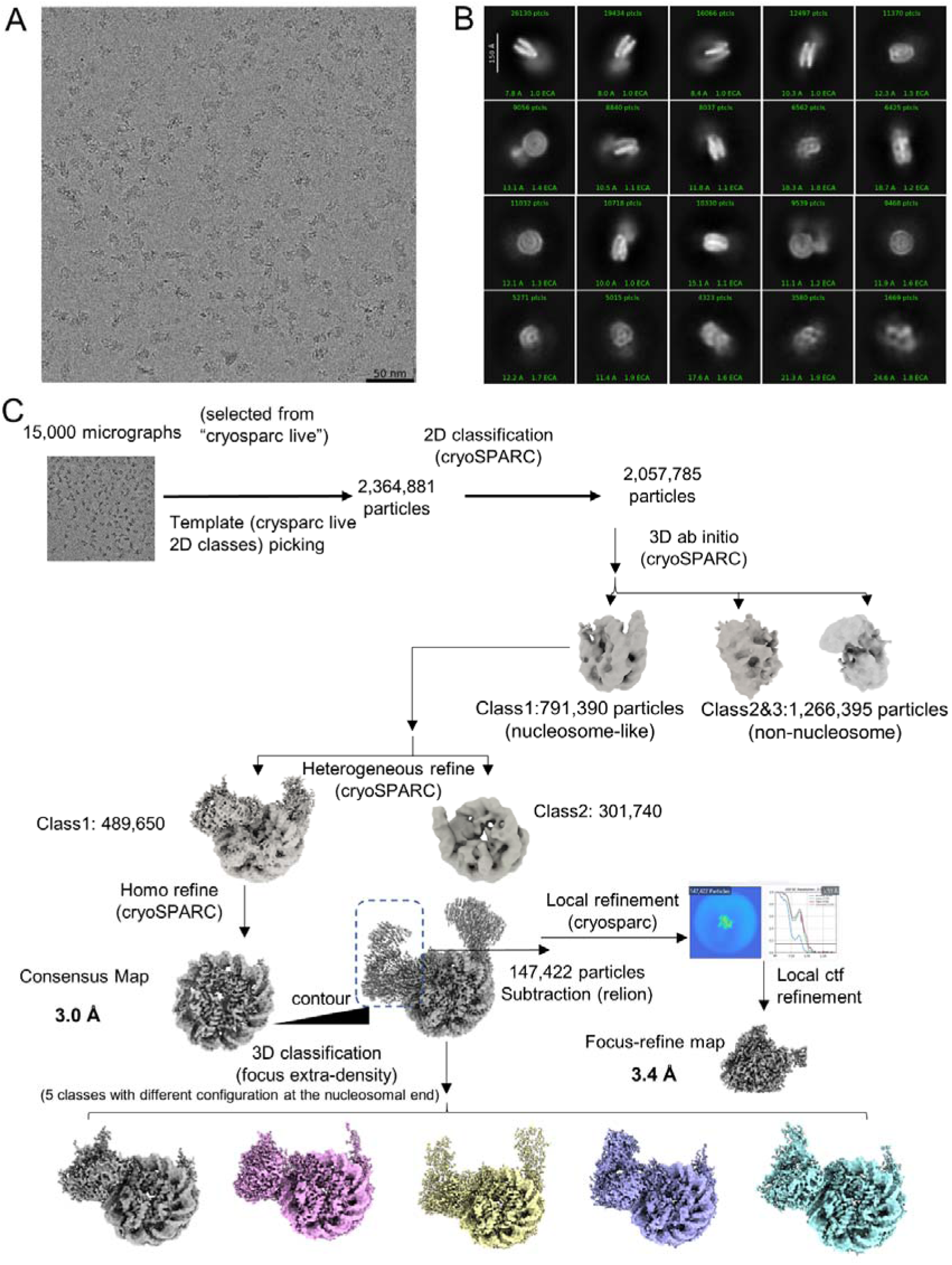
Cryo-EM data analysis. A. Representative micrograph. B. 2D class averages generated from the dataset obtained from CryoSPARC 4.6. C. Data processing strategy flow chart.

**Figure S4:**
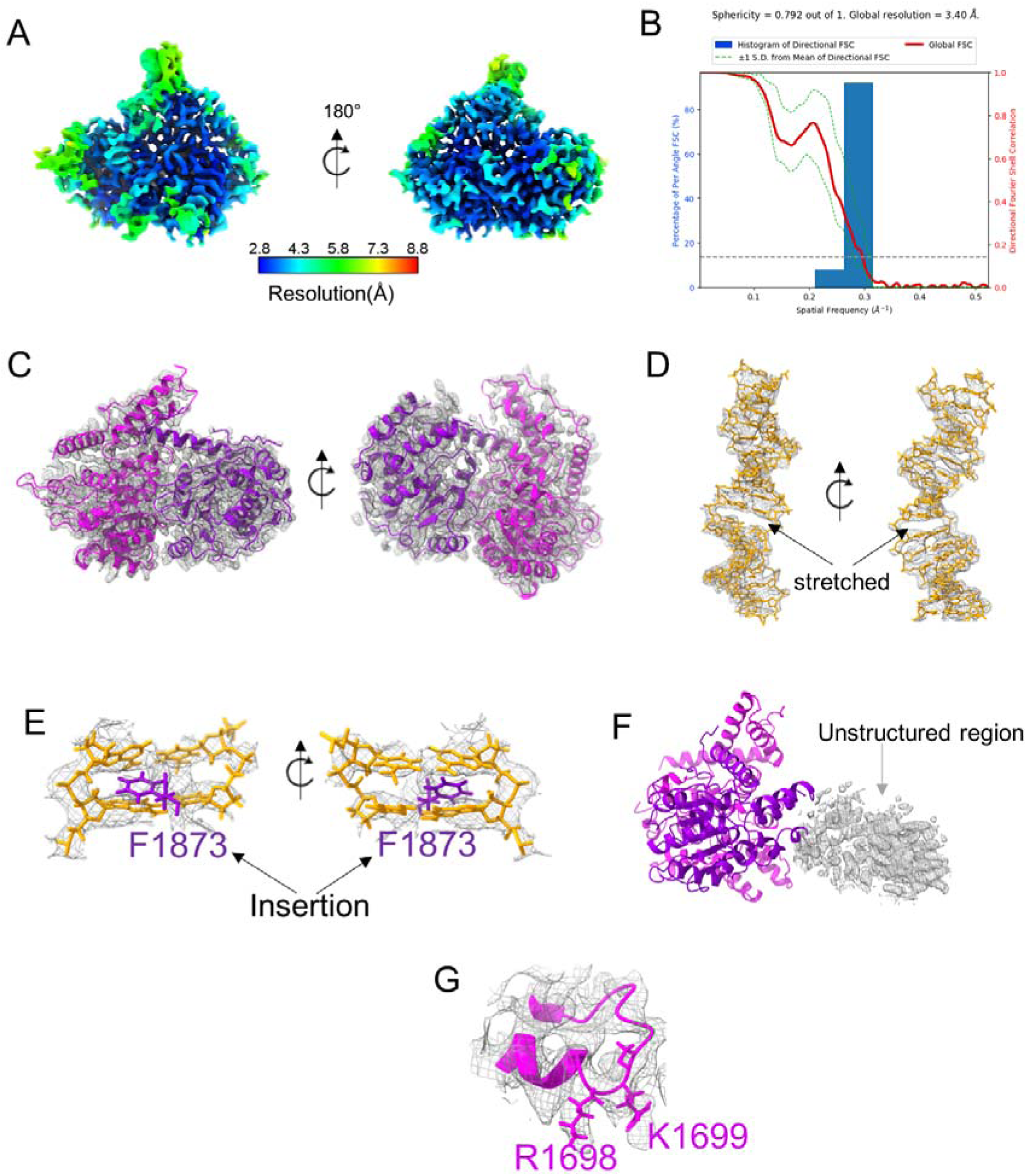
Cryo-EM data validation. A. Local resolution map of ATRX with the bound DNA (focus-refined from the consensus map by subtraction as described in the method). B. FSC curve of EM density in A. C-F: Representatives of model-to-density fitting for A.

**Figure S5:**
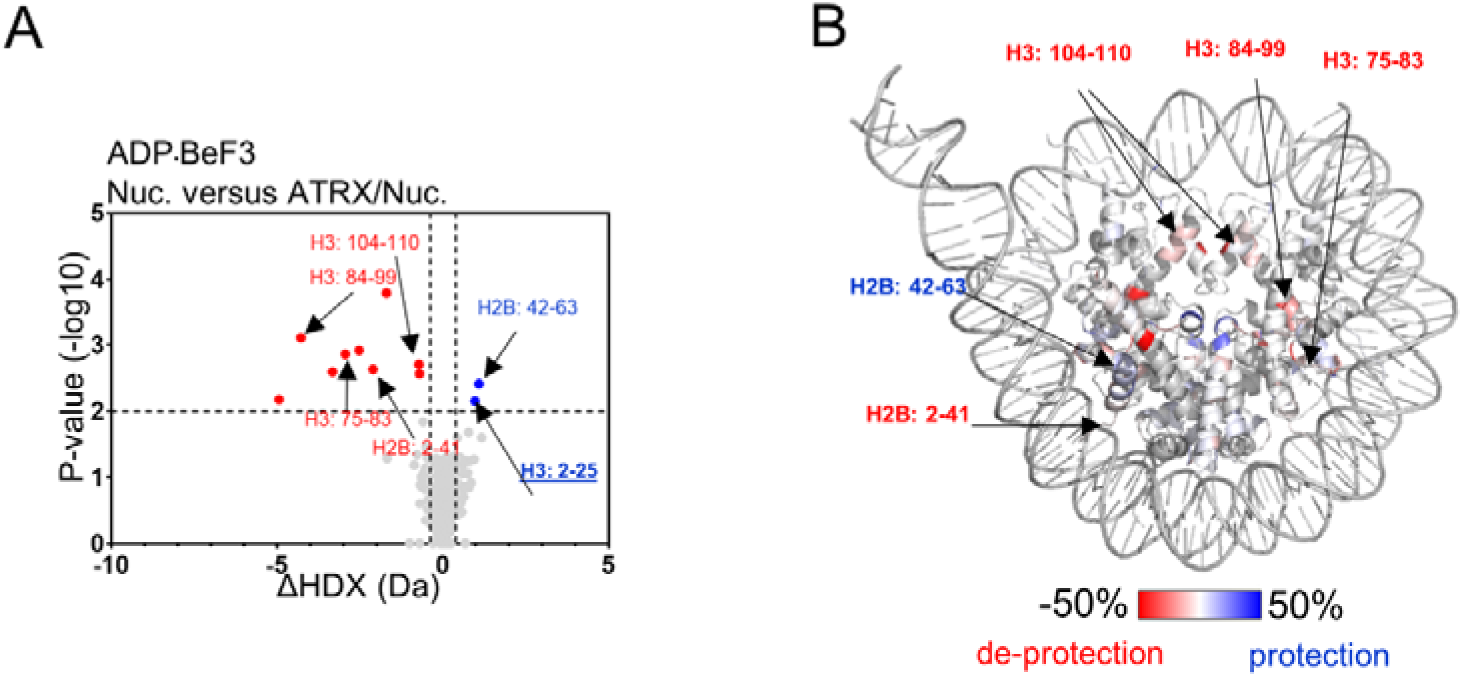
Deuterium uptake differences in HDX-MS for the nucleosome in complex with ATRX-C. The HDX difference between the nucleosome (histones) and the nucleosome with the “ADP•BeF3-bound” ATRX-C. The volcano plot in A and the colored structure model in B were generated in the same way as in Figure 1. Peptides showing significant changes were highlighted and labeled in both the volcano plot and the nucleosome structure model. The peptide on the H3 N-terminus is underlined. The significant cut-offs of Δ average |HDX|>0.383 Da and p-value<0.01 from Welch’s t-test are indicated by dashed lines.

**Figure S6:**
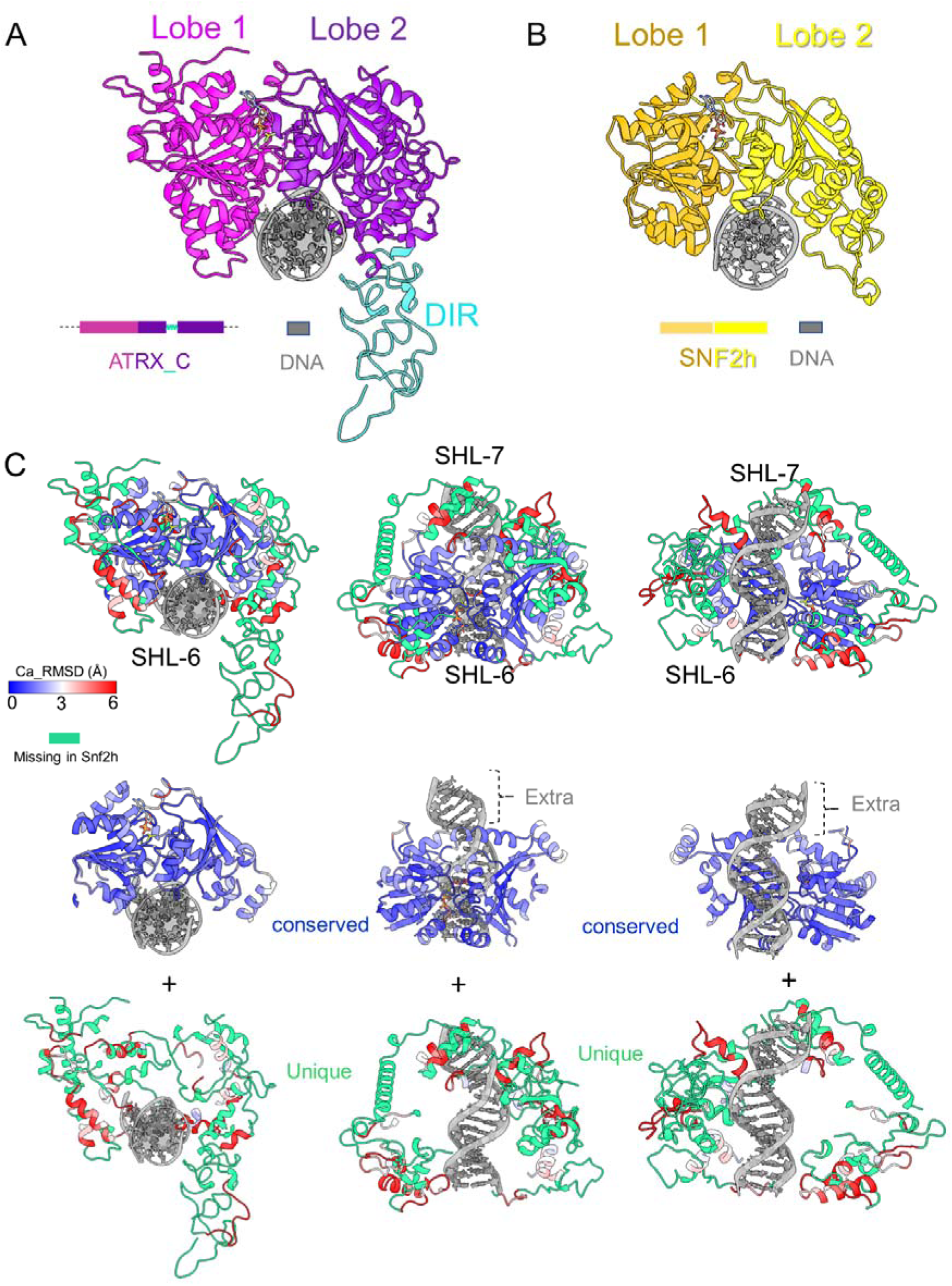
Structure comparison between ATRX^ATPase^ and Snf2h^ATase^. A. Structural model of ATRX^ATPase^. The color scheme for the domains of ATRX and DNA was indicated. B. Structural model of Snf2h^ATPase^ (from PDB: 8v4y). The color scheme for the domains of Snf2h and DNA was indicated. C. Ca_RMSD analysis between ATRX^ATPase^ and Snf2h^ATPase^. The analysis was done by the “matchmaker” command in chimeraX. The blue-white-red scale was used to describe the ca_RMSD value (from 0 to 6). The green color highlights regions of ATRX that are missing in Snf2h (which exists only in ATRX).

**Figure S7:**
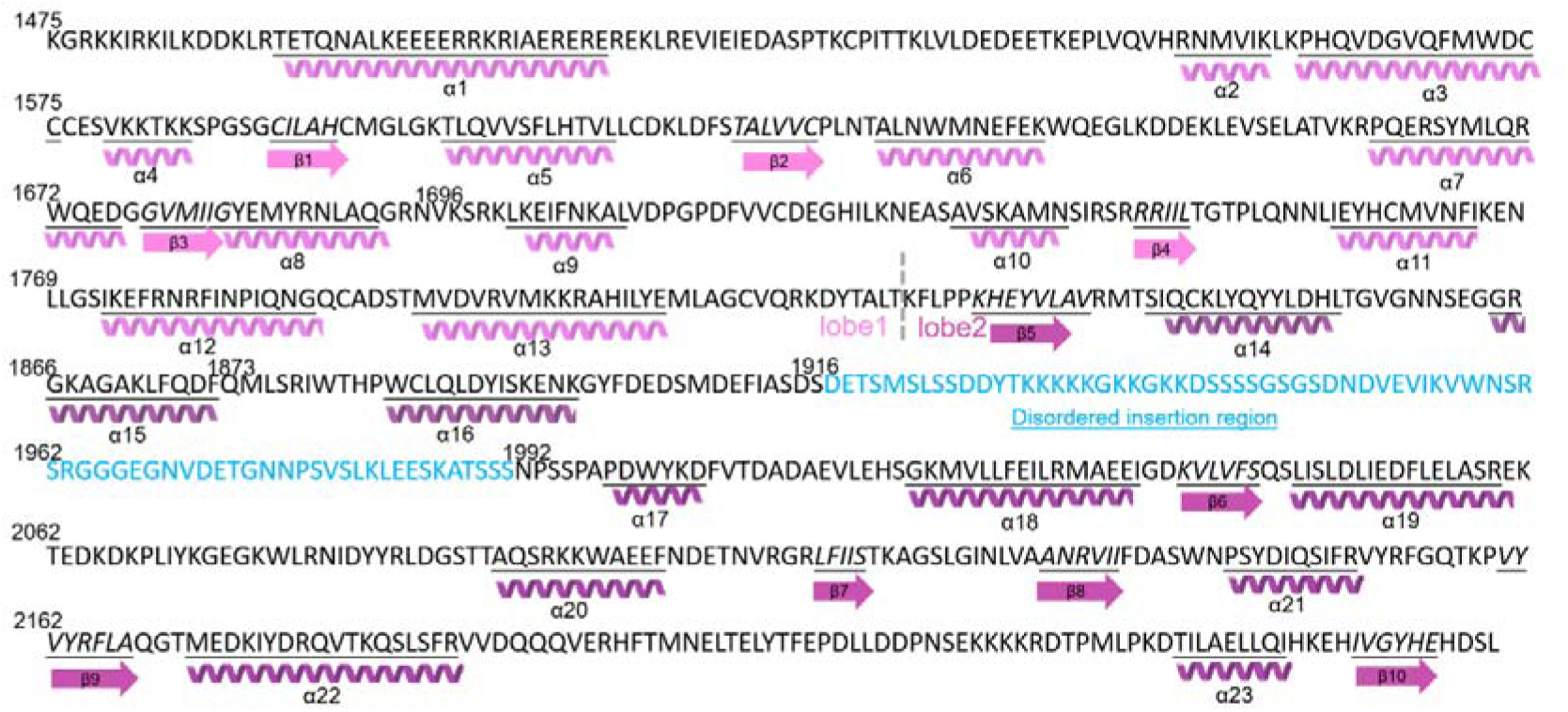
Sequence analysis of ATRX-C structure. Secondary structure elements revealed by cryo-EM are labeled with the sequence. “□” stands for □-helix, “β” stands for β-sheet.

**Figure S8:**
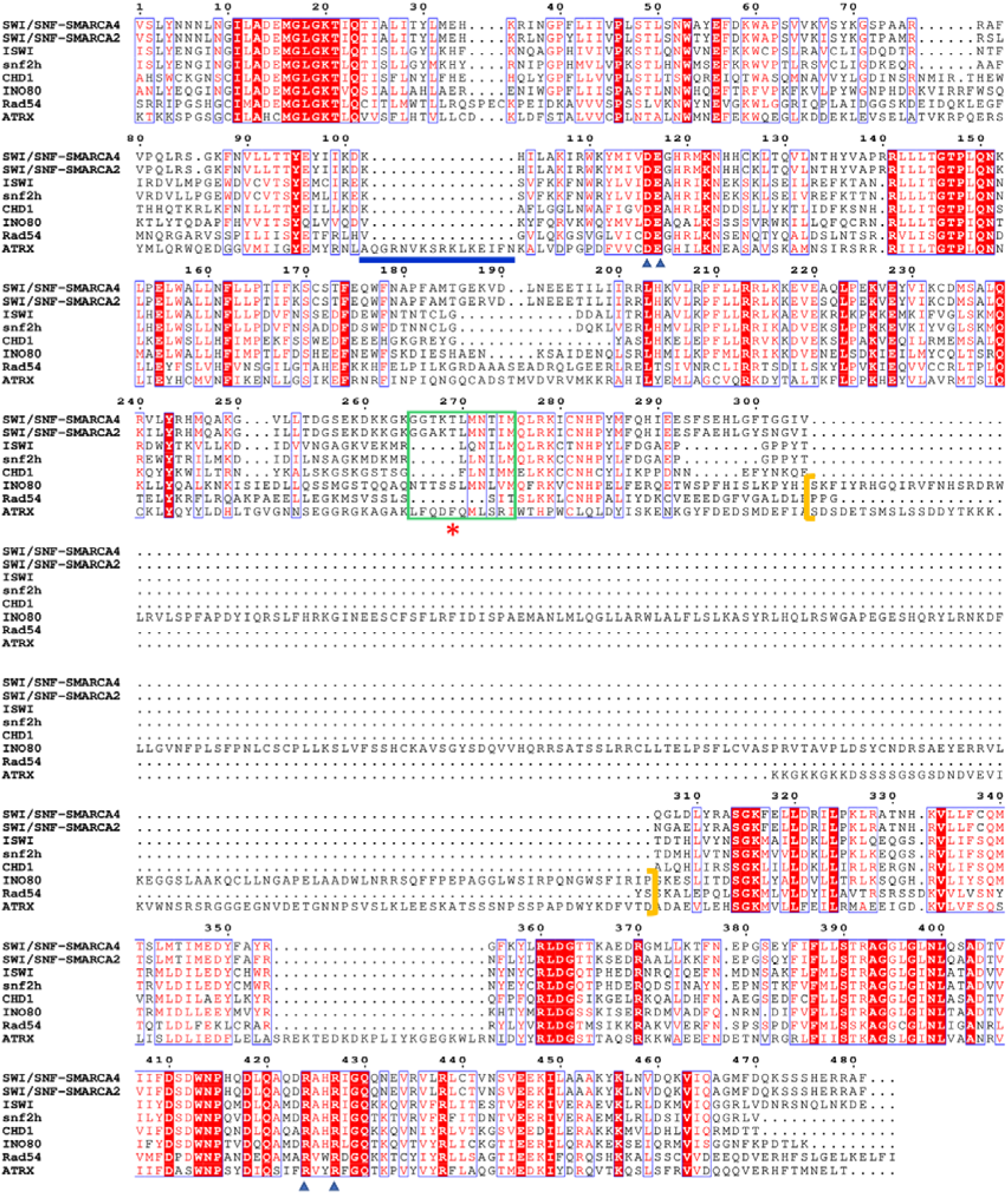
Sequence alignment of ATPases from ATRX and other remodelers. The sequence alignment focuses on the ATPase domains of ATRX, SWI/SNF (SMARCA2 and SMARCA4), ISWI, snf2h, CHD1, INO80, and Rad54. The green box highlighted the gating helix, red * highlighted the pin element F1873 in ATRX, the blue line highlighted the extra loop in ATRX for the “cross-gyre” interaction, and the brown square brackets indicated the disordered insertion region DIR.

**Figure S9:**
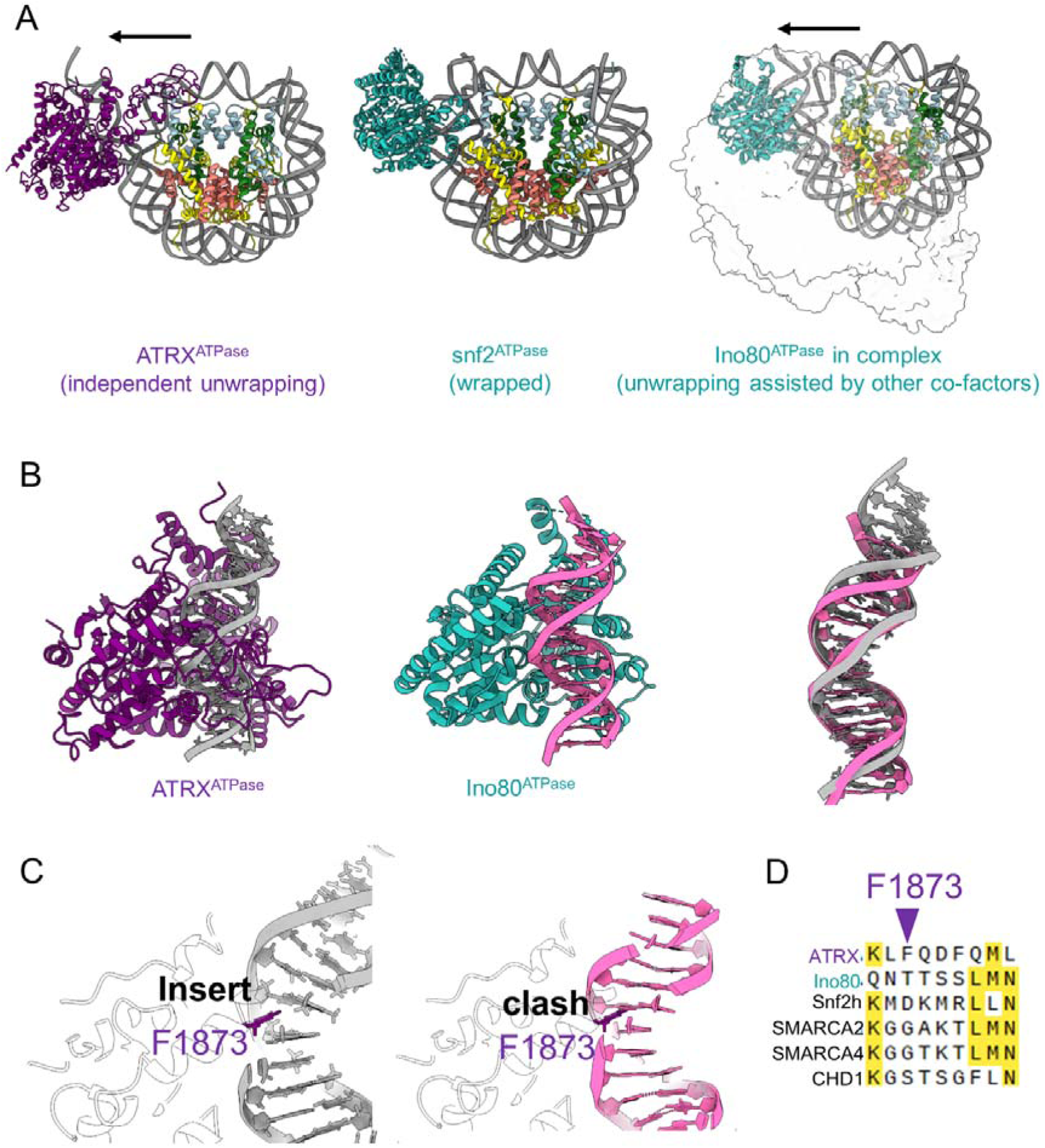
Comparison of ATPase-DNA interactions between ATRX and Ino80. A. ATRX^ATPase^ unwraps the DNA independent from co-factors. The structures of ATRX/nucleosome, Snf2^ATPase^/nucleosome at SHL6 (PDB 5×0×), and Ino80/nucleosome (8atf) were presented in parallel. B. ATRX^ATPase^ deforms DNA more profoundly than the Ino80 complex. The stretched minor groove by ATRX is even wider than the one in the Ino80 complex. C. The phenylalanine (F1873) inserted between the bases in the ATRX structure would clash with the DNA base in the Ino80 structure. D. Phenylalanine (F1873) is unique to ATRX among chromatin remodelers.

**Figure S10.**
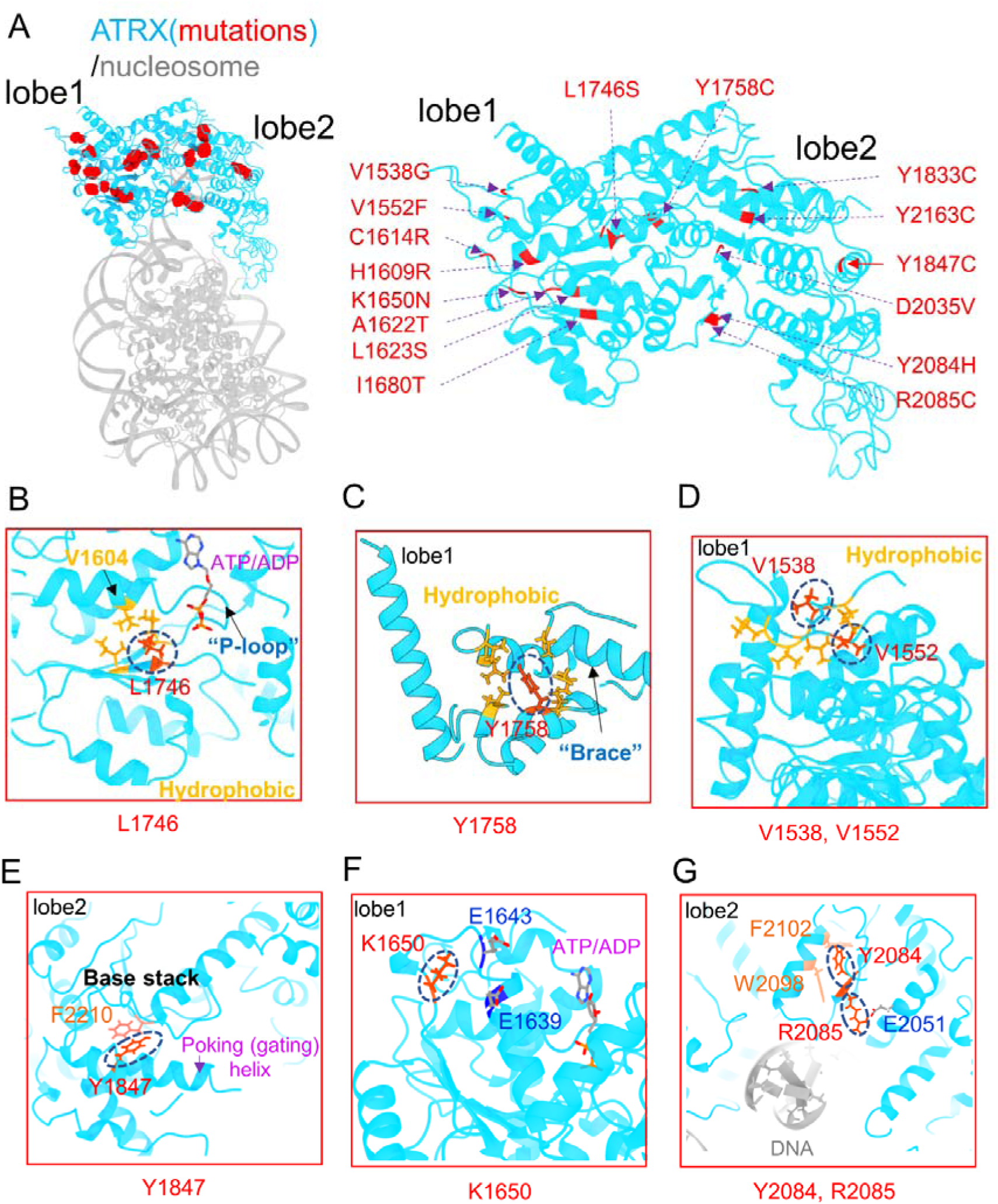
Mutations of ATRX^ATPase^ in reported diseases. A. The reported mutation sites^36,37^ were highlighted in red (ATRX ATPase is colored in cyan, and the nucleosome is colored in gray). Right: The detailed mutations were mapped in the ATPase. B-G. The local structure of the representative mutation sites.

## STAR Methods

### Resource Availability

#### Materials availability

Histone expression constructs, ATRX expression constructs, as well as vectors are available upon request.

#### Data and Code Availability

Coordinates and density maps will be deposited in the PDB (9L06) and EMDB databases (EMD-62684, EMD-62686, EMD-62690).

### Purification of human ATRX proteins from Sf9 insect cell

Human ATRX, ATRX-C (1251-2492), and ATRX-C mutations ΔDIR (Δ1916-1995), KSRK mutant (1696-1699: KSRK to ASAA), F1873A were expressed in the Bac-to-Bac system and purified from Sf9 insect cells, respectively. Cells were collected at 4000 r.p.m. (Eppendorf, centrifuge 5430 R, Rotor F-35-6-30) for 5 min, and lysed by Emerson SFX 550 (On 1 s, off 1 s, total 1 min) at 4 °C, in 600 mM NaCl, 20 mM Tris-HCl (pH 7.5), 10 mM imidazole, 1 pcs per 40 mL Protease Inhibitor Cocktail mini-Tablet (MCE, Cat. No.: HY-K0011), 10 μL per 40 mL DNase I (NEB, Catalog # M0303L), 1 mM MgCl2, 1 mM CaCl2. The cell lysate was spun down at a speed of 16000 r.p.m. for 30 min by Thermo Sorvall Lynx 6000 with rotor A-27. The supernatant was collected and then loaded into a gravity column with nickel agarose beads (GOLDBIO, Catalog ID H-320). After being eluted by 5 mL 500 mM imidazole, 150 mM NaCl, and 20 mM Tris-HCl (pH 7.5), the His-tagged protein was purified by a prepacked 5-ml HiTrap Q HP column. Protein was concentrated to 10-20 μM and stored in 400 mM NaCl, 20 mM Hepes (pH 7.5), 0.5 mM EDTA, 1 mM DTT, 10% glycerol. For the de-phosphorylation of ATRX, the protein was treated by CIP (1000 U CIP per mg protein) at RT for 1 hour after the Q HP step, and then applied to the size exclusion column in 300 mM NaCl, 25 mM Hepes (pH 7.5). Protein quality was analyzed by 4-15% SDS-PAGE. Protein was then stored with 10% glycerol. All purification columns were purchased from GE Healthcare.

### Plasmid construction for mutated ATRX-C

Specific primers were designed according to requirements (substitution or internal deletion) and used in the PCR reaction system with Phusion® High-Fidelity DNA Polymerase (NEB, M0530L). After amplification, 2 μL DpnI (NEB, R0176L) and 10 μL rCutSmart™ Buffer were added to 100 μL PCR reaction system, and then incubated at 37 °C for 1 h to remove template DNA. The PCR product was purified by TaKaRa MiniBEST DNA Fragment Purification Kit Ver.4.0 (Cat. # 9761), and then transformed into DH5α competent cell. After recovery overnight, bacteria were applied on LB agarose plate with ampicillin for growth. After culture and sequencing, colonies with the correct DNA sequence were stored.

### Nucleosome reconstitution and ATRX binding assay

Recombinant human histones H3.3-H4 and H2A-H2B were expressed and purified as described^38^. Purified H2A-H2B, H3.3-H4, and 207bp DNA (30bp flanking DNA with 147bp “widom 601” sequence^39^ at the center) were mixed at a ratio of 2.4: 2.4:1.0. Nucleosome was reconstituted by gradient dialysis^38^, and the products were analyzed by 5% native-PAGE.

The de-phosphorylated ATRX protein is mixed with reconstituted nucleosomes (250 nM) and titrated at 1:1, 2:1, 4:1 and 8:1 in the buffer of 20mM Tris (pH 8.0), 100mM NaCl and 1 mM TCEP. The binding assay was performed in apo state, with ADP (0.5 mM ADP, 1 mM MgCl2) or with ATP analogue (0.5 mM ADP, 1 mM MgCl2, 8 mM NaF, 1 mM BeSO4). The binding product/complex was then analyzed by 5% native-PAGE and 4-15% SDS-PAGE.

### MNase digestion assay

800 nM ATRX-C was mixed with 100 nM nucleosome (207 bp DNA, H3.3-H4), and incubated at room temperature for 10 min. Increasing amounts of MNase were added to each group and incubated at 37 °C for another 10 min. To quench the reaction, 3 μL of 0.5 M EDTA was added. DNA fragments were then purified by MinElute PCR Purification kit (QIAGEN), qualified through native gel and quantified via Agilent 2100 Bioanalyzer as described before^40^. ATP analogue (0.5 mM ADP, 1 mM MgCl2, 8 mM NaF, 1 mM BeSO4) or ADP (0.5 mM ADP, 1 mM MgCl2) was added to determine their influence on the protection function provided by ATRX-C to nucleosomal DNA.

### Nucleosome mobilization/sliding assay

Nucleosome mobilization activity was measured as previously described^30^. Nucleosome was first reconstituted on an asymmetric DNA. In the reaction, 10 nM nucleosome was used, and 1 μM protein (same concentration for ATRX-full length, ATRX-C, and all mutated proteins) was then added, in buffer containing 20 mM Tris-HCl (pH 7.5), 50 mM KCl, 0.1 mg/ml bovine serum albumin, 1 mM ATP, 5 mM MgCl2, 1 mM DTT. Reactions were stopped with 150 ng DNA and then resolved on 8% native PAGE. Gels were quantified by ImageQuant™ TL analysis software.

### Glycerol gradient and gradient fixiation (GraFix) crosslinking

To form the gradient, 2 ml top solution (25 mM HEPES pH 7.5, 50 mM NaCl, 0.5 mM EDTA, and 10% glycerol) was first added to a tube. 2 mL bottom solution (25 mM HEPES pH 7.5, 50 mM NaCl, 0.5 mM EDTA, and 30% glycerol) was then added to the bottom of the tube (Beckman). The tube was placed on a gradient maker (Biocomp 108) to form a continuous gradient. 200 ul of the sample at 5 μM was slowly loaded on top of the solution and spun at 4°C for 6 h at 60,000 rpm (Thermo Scientific WX100+, Rotor TH-660). The correct sample fractions were dialyzed against 50 mM Tris-HCl pH 7.5, 50 mM NaCl, 0.5 mM EDTA, and 1 mM DTT to remove the glycerol.

For GraFix, the same procedure was applied with the addition of 0.15% glutaraldehyde in the bottom solution to form the continuous crosslinker gradient. After GraFix, the samples (ATRX-C-nucleosome with or without ADP/ADP-BeFx) were fractionated and dialyzed against 50 mM Tris-HCl pH 7.5, 50 mM NaCl, 0.5 mM EDTA, and 1 mM DTT to quench the crosslinking reaction and remove glycerol.

### Single-particle cryogenic electron microscopy (cryo-EM) data processing

The crosslinked complexes of ATRX-nucleosome (Apo, ADP or ADP-BeFx) were concentrated to 5 uM. UltrAuFoil grids (R 1.2/1.3, Au 300) were glow discharged (EMItec, Lohmar, DE) at 15 mA for 1 min, and 4 ul sample was loaded onto the grid before plunge freezing by Vitrobot. Images for the complexes were acquired at a nominal magnification of 130000x on Krios G4 Cryo-TEM (300 kV), equipped with a Falcon 4i Direct Electron Detector (Pixel size was 0.9557Å). The movies were captured with an electron dose rate of 8 electrons per pixel per second for 5.0 s in EER format. The defocus range was _0.8 to _2.0 um.

The raw data was processed (motion correction and CTF estimation) by cryoSPARC (v4.6.0)^41^. Images were evaluated by inspecting the CTF fitting resolution. ∼15000 micrographs higher than 10 Å were selected. Roughly 1000 particles were manually picked for generating auto-picking templates through 2D classification in CryoSPARC. After several rounds of 2D classification to discard bad particles, 2,057,785 particles were selected for the ab initio classification. The particles in the nucleosome-like class was selected and further processed by heterogeneous refinement in cryoSPARC with the mask focusing on the extra density on the nucleosome. This separated nucleosome particles and complex particles. The complex 3D volume was further refined by the homogenous refinement as a consensus map. The aligned particles in the consensus map were transferred to the relion (v4.0)^42^ through pyem (https://github.com/asarnow/pyem). The particles were further classified without alignment (T value=20). The class with the clear extra density (∼147,422 particles) was refined in relion. The mask focusing the extra density was generated for subtraction. The initial map of the subtraction was reconstructed in relion. The particles were then transferred to the cryoSPARC for the local refinement, followed by the local ctf refinement. The final map was evaluated by the “local resolution estimation” in cryoSPARC and the 3DFSC server^43^.

### Modeling

The alphafold3 generated the Initial model of ATRX^44^. The initial model of the nucleosome was based on the PDB: 8v4y. The unwrapped 18 bp DNA end was generated by PyMol command “fnab”. All histones and DNA sequences were corrected by COOT^45^. The composite density map (ATRX density aligned with the consensus map) was used for the model assembly. The assembled model was then refined by ISOLDE^46^. Phenix real-space refinement was performed for the final model^47^.

### HDX-MS

15 μM ATRX-C was prepared in 150 mM NaCl, 20 mM HEPES pH 7.5 at 25 °C, 10 μM 207H3.3 nucleosome was prepared in 50 mM NaCl, 20 mM Tris-HCl pH 7.5 at 25 °C, 1 mM EDTA, 1 mM DTT. For the complex, the molar ratio between ATRX-C and nucleosome was 1.5:1. HDX was initiated by diluting the samples with the same buffer as ATRX-C, prepared in D2O (99.9%, Cambridge Isotope Laboratories) in a 1:9 ratio (90% D, pH 7.5 at 25 °C). The control experiment was conducted according to the procedures above, but replacing the D2O ATRX-C buffer with the H2O buffer. After 60 s, 600 s, or 3600 s of deuteration, the reaction mixtures were quenched by the addition of a quenching solution (1% formic acid, 3.84 M guanidinium chloride) in a 1:1 (v/v) ratio at 0.5 °C to lower the pH of the mixture to approximately 2.5. The quenched mixtures were injected through a nepenthesin-2/pepsin column (AffiPro, 2.1 mm × 20 mm) at room temperature for online digestion, followed by desalting in a C18 cartridge (ThermoFisher, 100 Å, 5 μm, 300 μm × 5 mm) at 0.5 °C for 3 minutes using a flow rate of 120 μL/min with 0.1% formic acid. The desalted peptides were then eluted and separated on a Hypersil GOLD C18 column (ThermoFisher, 175 Å, 1.9 μm, 50 mm × 1 mm) at 0.5 °C with a solvent gradient of 0.1% formic acid and acetonitrile containing 0.1% formic acid as solvents A and B, respectively. The flow rate was set to 45 μL/min, with the gradient as follows: 0 to 0.5 min: 5% to 8% B, 0.5 to 12 min: 8% to 40% B, 12 to 13.5 min: 40% to 95% B, 13.5 to 15.5 min: 95% B, 15.5 to 15.6 min: 95% to 5% B, 15.6 to 17 min: 5% B. All the deuterium exchange steps, online digestion and liquid chromatography parts were done automatically via LEAP HDX automation (Trajan) connected with an UltiMate 3000 RSLCnano pump module (ThermoFisher). Eluted peptides were introduced directly into an Orbitrap Fusion Lumos Mass Spectrometer (ThermoFisher). The global parameters were ion source type, H-ESI; spray voltage, static; positive ion (V), 3600; ion transfer tube temperature (°C), 275; vaporizer temperature (°C), 100. Mass spectra was acquired under 60000 resolution in the MS Orbitrap step. Peptide sequences were identified using Proteome Discoverer 2.5 (ThermoFisher), and the relative deuterium uptake for each peptide were analysed by HDExaminer (Sierra Analytics Inc.). Volcano plots were generated by GraphPad Prism 9 software; the significant cut-offs of average HDX change are generated by multiply repeatability by 2.776 for each state under p<0.05. Fractional uptake or fractional uptake differences were mapped to the protein structures in a blue-white-red color scale by PyMOL.

### NADH-coupled ATP hydrolysis assay

ATP Hydrolysis assay was performed in hydrolysis buffer that contains 50 mM HEPES pH 7.5, 100 mM KCl, 4 mM MgCl2, 0.7 mM NADH, 1 mM phosphoenolpyruvate, 1 mM DTT and 2.4 μL (per 100 μL) pyruvate kinase-lactate dehydrogenase. For 1:1 ratio of protein to activator in the reaction, 100 nM protein and 100 nM activator (free DNA or Nucleosome) were mixed. The addition of 1 mM ATP was used to initiate the reaction. The absorbance of NADH was measured at 340 nm wavelength over 180 minutes (recording every 20 minutes). To convert the raw data to ATP hydrolysis rate ([M]/s), the A340/s raw readings were divided by the NADH extinction coefficient (6330 M−1 cm−1). Background ATP hydrolysis (ATRX protein without any activator or blank buffer without protein and activator) was subtracted, and then rates were calculated and determined by fitting to a linear regression^48^.

